# Scalable Species Tree Inference with External Constraints

**DOI:** 10.1101/2021.11.05.467436

**Authors:** Baqiao Liu, Tandy Warnow

## Abstract

Species tree inference under the multi-species coalescent (MSC) model is a basic step in biological discovery. Despite the developments in recent years of methods that are proven statistically consistent and that have high accuracy, large datasets create computational challenges. Although there is generally some information available about the species trees that could be used to speed up the estimation, only one method–ASTRAL-J, a recent development in the ASTRAL family of methods–is able to use this information. Here we describe two new methods, NJst-J and FASTRAL-J, that can estimate the species tree given partial knowledge of the species tree in the form of a non-binary unrooted constraint tree.. We show that both NJst-J and FASTRAL-J are much faster than ASTRAL-J and we prove that all three methods are statistically consistent under the multi-species coalescent model subject to this constraint. Our extensive simulation study shows that both FASTRAL-J and NJst-J provide advantages over ASTRAL-J: both are faster (and NJst-J is particularly fast), and FASTRAL-J is generally at least as accurate as ASTRAL-J. An analysis of the Avian Phylogenomics project dataset with 48 species and 14,446 genes presents additional evidence of the value of FASTRAL-J over ASTRAL-J (and both over ASTRAL), with dramatic reductions in running time (20 hours for default ASTRAL, and minutes or seconds for ASTRAL-J and FASTRAL-J, respectively).

**Availability:** FASTRAL-J and NJst-J are available in open source form at https://github.com/RuneBlaze/FASTRAL-constrained and https://github.com/RuneBlaze/NJst-constrained. Locations of the datasets used in this study and detailed commands needed to reproduce the study are provided in the supplementary materials at http://tandy.cs.illinois.edu/baqiao-suppl.pdf.

## 1 Introduction

Species tree inference is a common task in phylogenetics and serves as a prior step for many downstream analysis (e.g., understanding adaptation). However, gene tree discordance, where the gene tree topology can differ from the species tree topology, presents unique challenges for species tree inference. Major sources of gene tree discordances include horizontal gene transfer (HGT), gene duplication and loss, and notably incomplete lineage sorting (ILS) [1]. ILS is modeled statistically by the multi-species coalescent (MSC) model [2].

A common approach to species tree estimation under the MSC model concatenates the alignments of genomic regions into a single “super-alignment” and then runs approaches such as maximum likelihood on the alignment. This “concatenation” analysis, however, can be statistically inconsistent [3] under the MSC. An increasingly popular alternative approach first infers trees from each genomic region and then applies a “summary method” to estimate the species tree from these “gene trees”. ASTRAL [4, 5, 6] is the most well known summary method and is statistically consistent under the MSC. ASTRAL runs in time that is polynomial in its input, and is often more accurate than concatenation analyses when the level of ILS is high. However, ASTRAL can also be computationally intensive when the number of species is large or when there is a large number of gene trees that are topologically very different from each other. Other statistically consistent summary methods include MP-EST [7], NJst [8], ASTRID [9], and FASTRAL [10], but ASTRAL is currently the main summary method used on large phylogenomic datasets.

In this paper we address the problem of estimating a species tree from a set of gene trees when there is partial knowledge of the species tree, given in the form of an incompletely resolved (i.e., multifurcating) constraint tree. This is a natural problem in phylogenetics, especially for those datasets (e.g., avian or mammalian phylogeny) that have received intense study over many years and yet remain controversial [11, 12, 13, 14, 15, 16, 17, 18]. A recent variant of ASTRAL [19], which we refer to as “ASTRAL-J”, was developed to allow the user to specify such a constraint tree, and then algorithmically ensures that the tree returned by ASTRAL-J will be a refinement of the constraint tree (i.e., the returned tree will resolve the polytomies in the constraint tree, and so will have additional edges).

Here we introduce two new methods, NJst-J and FASTRAL-J, that are designed to estimate species trees from a set of gene trees, given a non-binary unrooted constraint tree. We prove that these methods are polynomial time and statistically consistent under the MSC provided that the given constraint tree has no false positive edges. Our simulation study shows our new methods are faster than ASTRAL-J and that the better of the two methods (FASTRAL-J) matches or improves on ASTRAL-J for accuracy. Finally, using a constraint tree based on the “Magnificent Seven” clades from Reddy et al. [13] applied to the Jarvis et al. [11] avian dataset with 48 species and 14,446 genes shows FASTRAL-J provides dramatic reductions in runtime over default ASTRAL (20 hours for default ASTRAL and 1.5 minutes for FASTRAL-J).

### 2 Background

**Basic definitions** All trees are assumed to be unrooted and on leafset |*S*|, with *S* = *n*. We also assume that gene trees evolve within a species tree under the multi-species coalescent (MSC) model. Each unrooted tree *T* can be represented by its set *C*(*T*) of bipartitions on its leafset produced by deleting edges in *T*. Tree *A* is said to be a **refinement** of tree *B* if ⊆ *C*(*B*) *C*(*A*). The Robinson-Foulds (RF) distance [20] between two trees *A* and *B* on *S* is defined to be the |*C*(*A*)Δ*C*(*B*) | (i.e., bipartitions in one but not the other tree), and the RF error rate is *RF* (*A, B*)*/*(2*n* − 6).

A **summary method** takes as input a set 𝒢 of unrooted gene trees on leafset *S* and returns an estimated species tree. We will say that summary method *f* is “tree-constrained” if the input includes a (potentially multifurcating) tree 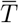 on *S*, and we constrain the output tree *T* to satisfy 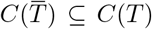. We reuse the nomenclature of [19] and refer to 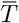 as the “backbone”. We will say the backbone is “valid” if it has no false positive edges (i.e., every bipartition in the backbone appears in the true species tree, and so the backbone is a contraction of the true species tree).

#### Definition 1.

*A tree-constrained summary method f is said to be* ***statistically consistent under the MSC given a valid backbone*** *if the probability that f returns the true species tree converges to* 1 *as the number of gene trees increases*.

### ASTRAL and *X*-constrained MQSST

We first define the maximum quartet-support supertree (MQSST) problem. The input is a set 𝒢 of unrooted binary gene trees, each on the same set *S* of leaves, and the output is an unrooted binary tree *T* on *S* that maximizes

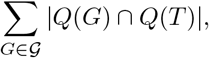

where *Q*(*t*) denotes the set of quartet trees of *t*. MQSST is NP-hard [21], but an exact solution to MQSST is a statistically consistent method for species tree estimation under the MSC [4].

ASTRAL solves the **X-constrained MQSST problem**, which we now define. The input is a set 𝒢 of gene trees and also a set *X* of “allowed bipartitions”, and the output is a tree *T* optimizing the MQSST score subject to *C*(*T*) ⊆ *X*. ASTRAL [4] uses dynamic programming (DP) to solve the *X*-constrained MQSST problem in polynomial time. Furthermore, by constructing *X* so as to contain all the bipartitions from the input gene trees, ASTRAL is guaranteed to be statistically consistent under the MSC [4].

### ASTRAL-J

ASTRAL-J [19] (named after the command line flag of this variant) allows the user to specify a constraint tree 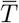 and then guarantees that the output tree *T* satisfy 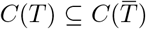. Given input set 𝒢 of gene trees and constraint tree 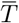, ASTRAL-J associates to every *G* ∈ 𝒢 a binary tree *t*_*G*_ that refines 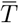. To construct *t*_*G*_ given binary gene tree *G* and 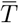, ASTRAL-J adapted a linear-time tree algorithm [22] referred to as B-RF(+) (itself an improvement upon OCTAL [23] in running time) into the B-RF(*) algorithm for this transformation of *X*. Concretely, the B-RF(*) algorithm returns a tree, referred to as *Comp*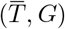, that has the minimum Robinson-Foulds (RF) distance to *G* among all refinements of 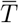. We informally call this step the “B-RF(*) transformation of trees in 𝒢”. As the B-RF(*) algorithm produces trees that may have polytomies, ASTRAL-J then employs heuristics from ASTRAL to produce binary refinements of these trees. The binary tree thus associated to *G* is referred to as *t*_*G*_, and by construction is a binary tree that refines 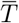.

ASTRAL-J also computes additional binary trees that are refinements of 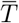; the goal of this step is to expand the search space (and hence potentially improve topological accuracy) while still enforcing the constraint tree 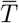. Finally, ASTRAL-J computes the set 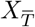 to be the bipartitions appearing in any of the produced trees (i.e., the trees *t*_*G*_ or the computed binary refinements of the consensus trees). ASTRAL-J then runs ASTRAL with input 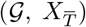 (i.e., the same set of gene trees but a set of allowed bipartitions that enforces compatibility of the output tree with 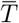). By design, therefore, ASTRAL-J guarantees that the output tree will be a refinement of 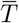.

### NJst and ASTRID

NJst [8] and ASTRID [9] are distance-based summary methods for estimating species trees under the MSC. Like ASTRAL, NJst and ASTRID are statistically consistent [24]. Each method uses the “internode distance matrix”, where for two leaves *x, y D*[*x, y*] is the average (across the gene trees) of the number of nodes on the path between *x* and *y*. Then NJst runs neighbor joining (NJ) [25] on the internode distance matrix while ASTRID runs balanced minimum evolution (BME) within FastME [26]. As proven in [24], under the MSC, as the number of genes increase the internode distance matrix converges to a matrix that is additive for the true species tree. Since NJ and BME within FastME both have a positive safety radius [27, 28], both NJst and ASTRID are statistically consistent under the MSC.

### FASTRAL

Although ASTRAL is polynomial time, its worst case running time grows almost quadratically in |*X*|, and by design, *X* contains all the bipartitions from all the input gene trees. Therefore, |*X*| will be large when the input gene trees are topologically very different from each other, which can be due to both high levels of ILS or gene tree estimation error. Hence, ASTRAL can be computationally intensive under a wide range of model conditions. FASTRAL speeds up ASTRAL by producing a smaller set of allowed bipartitions that nevertheless suffices to guarantee statistical consistency [10]. Here we describe how FASTRAL was used in [10], given the input set of *k* genes 𝒢:

1. Compute *Y* subsamples (with replacement) of, where at least one subsample is proportional to *k* in size
2. For each of the *Y* subsamples, run ASTRID on the subsample, thus producing *Y* ASTRID trees.
3. Let *X*′ denote the set of bipartitions in these *Y* ASTRID trees.
4. FASTRAL then invokes ASTRAL as a subroutine to solve the constrained-MQSST problem on (𝒢, *X*′) and outputs the result.

As proven in [10], FASTRAL is statistically consistent under the MSC. The specific settings for the subsamples explored in [10] used *Y* = 51, with one subsample containing all the genes, 10 subsamples containing half the genes, 20 subsamples containing 25% of the genes, and 20 subsamples containing 10% of the genes. This approach provided very good accuracy and improved the runtime substantially compared to ASTRAL.

## 3 New methods: NJst-J and FASTRAL-J

**NJst-J** Recall that NJst has two steps: first it computes the internode distance matrix and then it runs neighbor joining (NJ). NJst-J modifies the second step so that the returned tree is a refinement of the backbone 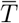. Recall that NJ uses an agglomerative clustering algorithm that begins with a star tree and then progressively makes pairs of nodes into siblings until the tree is fully resolved. Our modification to NJst uses the backbone 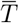 to determine whether to accept a siblinghood proposal. Effectively, in NJst-J we reject a siblinghood proposal if accepting the proposal would produce a tree that is incompatible with 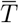. By construction, NJst-J will always return a tree, and the tree it returns will never violate the constraint tree.

### FASTRAL-J

As FASTRAL invokes ASTRAL as a subroutine to solve the DP problem on (𝒢, *X*′), we can reuse the construction from ASTRAL-J, and transform *X* to 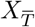 with respect to the backbone 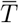 analogously. FASTRAL-J directly builds on FASTRAL and takes an additional input, the backbone tree 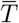; see Figure 1. FASTRAL-J executes steps 1 to 3 the same way as FASTRAL for constructing set *X*′. Then, FASTRAL-J constructs 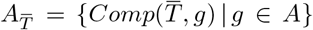, and resolves polytomies in 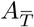 using FASTRAL’s heuristic for refining polytomies. Let 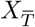 be the induced set of bipartitions of 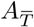. FASTRAL-J invokes ASTRAL to solve the constrained-MQSST problem on 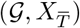.

**Figure 1:**
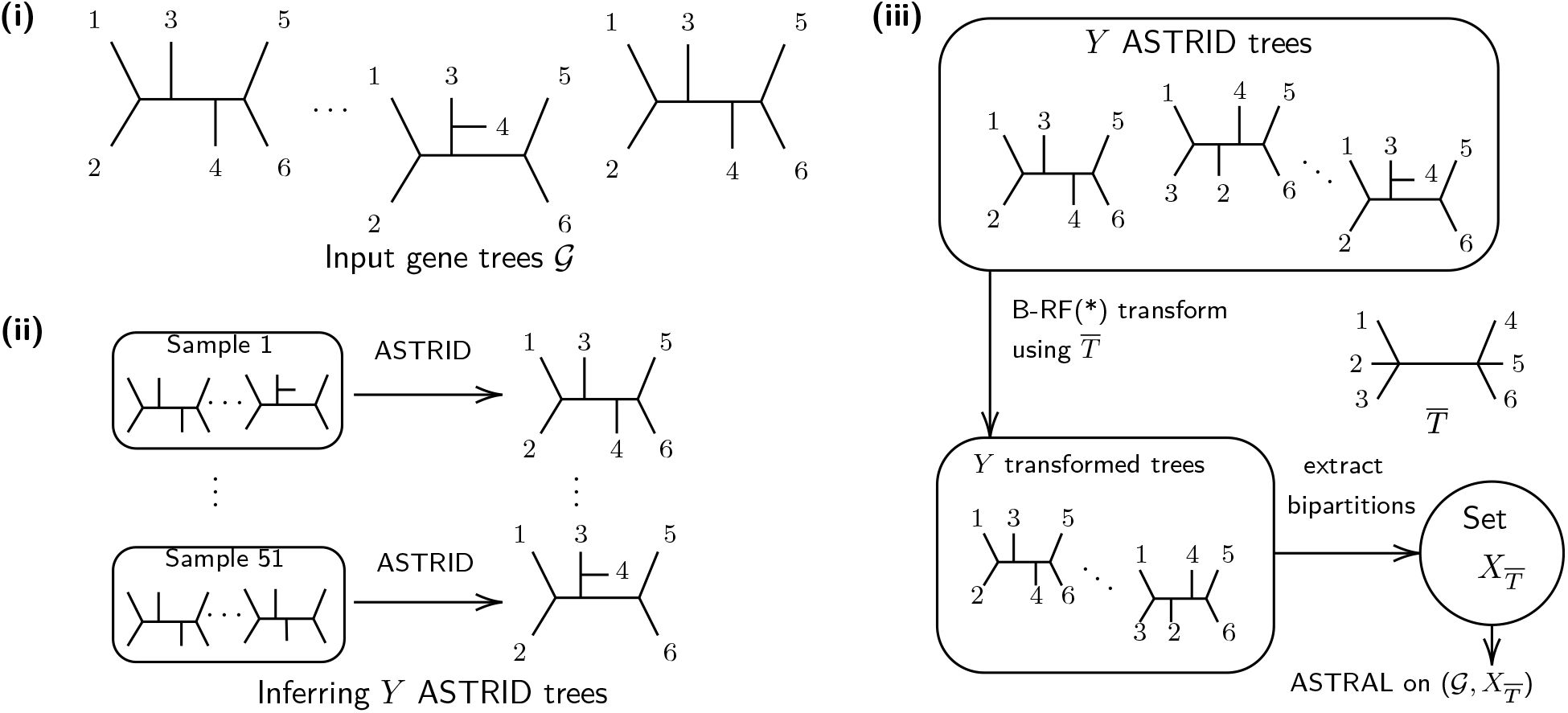
The FASTRAL-J pipeline. *(i)* The input unrooted gene trees. *(ii)* FASTRAL-J creates in total Y samples from 𝒢, and infers ASTRID species trees from these subsamples. *(iii)* The Y ASTRID trees are transformed using the B-RF(*) algorithm (developed in ASTRAL-J) to force them to become refinements of the user-defined constraint tree (the backbone 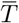, which here is 123|456). The polytomies of this transformed set of ASTRID trees are then resolved, and the bipartitions of these trees induce the search space set 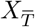. Finally ASTRAL is run on 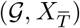.

## 4 Theoretical results

### Theorem 1.

*FASTRAL-J is statistically consistent under the MSC given valid backbones. Furthermore, if a polynomial number of ASTRID trees are computed, then FASTRAL-J runs in polynomial time*.

*Proof*. Let *T* be the unrooted topology of the true species tree and let 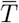 denote the valid backbone; hence, *T* refines 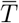. The proof that FASTRAL is statistically consistent under the MSC given in [10] operates by establishing that at least one of the ASTRID trees computed on the different samples will produce *T* with probability converging to 1 as the number of genes increases; the rest follows since the optimal solution to MQSST will converge uniquely to *T* as the number of genes increases (from [4]). The argument we provide here is very similar.

It suffices to show that the true species tree topology *T* is feasible in the limit. By construction, at least one subsampled gene set increases in size as the number of genes increase, and these samples are drawn at random from the set of possible genes. Hence, at least one produced ASTRID will converge to the true species tree *T* as the number of genes increase. Since *T* refines 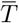, in the limit that ASTRID tree will refine 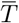 *T* with probability converging to 1. Note that among all trees that refine 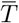, *T* has the minimum RF distance to itself. Therefore *T* ‘s bipartitions will appear in the search space of ASTRAL 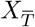 that is given by FASTRAL-J. As a result, *T* is a feasible solution to the MQSST problem that ASTRAL will run. Once the number of genes is large enough that the most frequently observed quartet tree on every four leaves is the most probable quartet tree for that set, the true species tree is the unique tree topology that optimizes the MQSST criterion. Hence, as the number of genes increases, FASTRAL-J will return the true species tree with probability converging to 1.

For the running time analysis, computing each ASTRID tree uses *O*(*kn*2) time to compute the internode distance matrix and then *O*(*n*3) to compute the FastME tree from the matrix. We only compute a polynomial number of ASTRID trees, so that the size of the constraint set given to ASTRAL is polynomial in *n*. Hence, the entire time is polynomial.□

### Theorem 2.

*NJst-J is polynomial time and statistically consistent under the MSC given valid backbones*.

*Proof*. We begin by reviewing the proof in [24] that NJst is statistically consistent under the MSC. The first part of this proof is that as the number of genes increases, the internode distance matrix *D* will converge to an additive matrix *A* for the species tree *T*, so that from a large enough number of genes the matrix *D* will satisfy *L*∞(*D, A*) *< f/*2 where *f* is the shortest length of any internal edge in the species tree *T*. The second part of the proof relies on the theorem by Atteson [27] that NJ will return the true species tree topology given such an estimated distance matrix *D* (i.e., one that is within *f/*2 of the species tree additive distance matrix); this is what is meant by saying that NJ has a positive safety radius, and is why NJst is statistically consistent under the MSC.

Now consider NJst-J, and assume that *L*∞(*D, A*) *< f/*2; in this case, the agglomerative clustering step within NJ will, in each iteration, only pick true sibling pairs of clusters. Therefore given an estimated distance matrix *D* satisfying *L*∞(*D, A*) *< f/*2, the modified NJ in NJst-J will behave exactly the same as the original NJ algorithm and so return the true species tree; in other words, NJst-J is statistically consistent under the MSC given a constraint tree without any false edges.

NJst-J uses *O*(*kn*2) to compute the internode distance matrix and then runs NJ (which is *O*(*n*3) time.

Hence, NJst-J runs in polynomial time.□

We end with a proof of consistency for ASTRAL-J.

### Theorem 3.

*ASTRAL-J is statistically consistent under the MSC given valid backbones*.

*Proof*. Let *T* denote the unrooted topology of the true species tree and 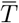 denote the valid backbone (thus, *T* refines 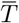). Theorem 2 of [4], under the MSC model, in the limit, *T* is the unique solution to the (unconstrained) MQSST problem. Therefore, it suffices to prove that in the limit (as the number of genes increases), *T* is in the feasible set of ASTRAL-J for this valid backbone. We first note that *T* has non-zero probability under the MSC, and so as the number of genes increases *T* will appear in 𝒢 with probability converging to 1. Recall that ASTRAL-J replaces its input gene trees *t* by other trees that are refinements of the backbone 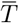 and that are the closest to *t* with respect to RF distance. However, when *t* is a refinement of the backbone, then there is no replacement needed. Because 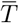 is a valid backbone, automatically *T* refines 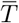. Hence, if *T* appears among the gene tree input, then the set of allowed bipartitions that ASTRAL-J produces will contain all the bipartitions of *T*. Hence, *T* will be a feasible solution to ASTRAL-J given the input set of gene trees and backbone 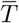.□

We note that although the above argument is straightforward, this result is not presented in [19].

## 5 Experimental Study

We evaluated our new methods, NJst-J and FASTRAL-J (using the same settings for FASTRAL as used in [10]) in comparison to ASTRAL-J (available at https://github.com/maryamrabiee/Constrained-search) on simulated datasets with 48-1001 species and 1000 genes, where ILS causes discordance between true gene trees and the species tree. NJst-J and FASTRAL-J are available in github (see abstract) and all datasets are available in public repositories. See http://tandy.cs.illinois.edu/baqiao-suppl.pdf for full details needed to reproduce the study.

For these datasets we had the true species tree (known to us since we performed a simulation study) and so could precisely quantify species tree estimation error. We also explored results on the avian phylogenomics dataset studied by Jarvis et al. [11] with 48 species and 14,446 genes. In both cases, we used gene trees estimated using maximum likelihood heuristics. We report the empirical statistics of these datasets in Table 1, showing number of taxa, number of genes, number of replicates per model condition, average discordance (AD) between true gene trees and true species tree, and gene tree estimation error (GTEE). For the AD and GTEE statistics, we report Robinson-Foulds (RF) error rates, which are RF distances divided by 2*n* − 6 (where *n* is the number of taxa) so as to produce a value between 0 and 1.

**Table 1:**
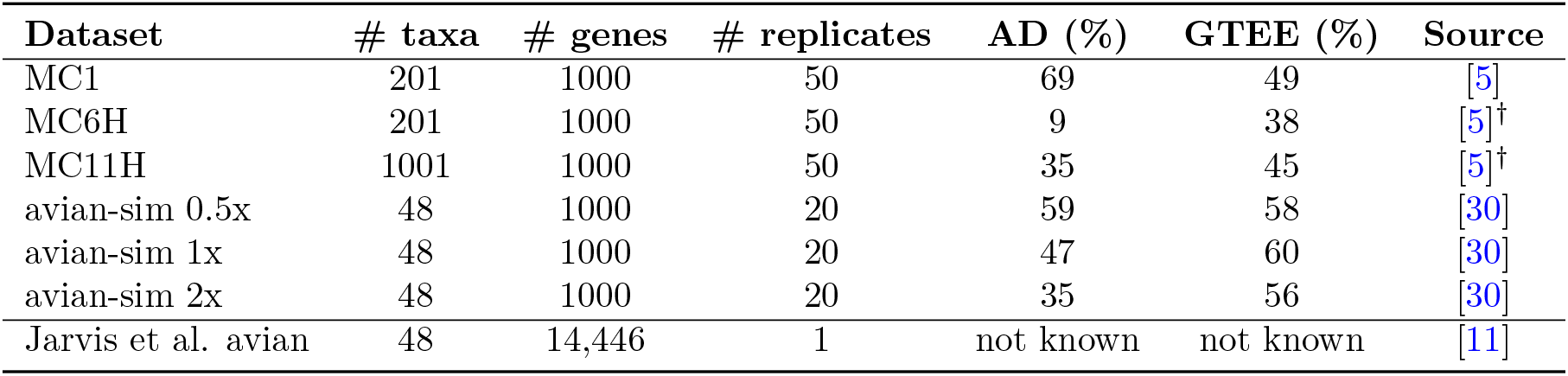
Empirical statistics for the datasets in this study (top datasets are simulated, bottom is biological). We report the number of taxa, number of gene trees, number of replicates, the degree of ILS as measured by AD (average discordance), and the amount of GTEE (gene tree estimation error). †: the MC6H and MC11H datasets are versions of MC6 and MC11 (from ASTRAL-II) with shorter alignments.

For the simulated datasets, we used the true species tree, and collapsed the branches that were short, as these are the branches that are statistically the hardest to reconstruct. We varied the percentage *B* of the branches that were retained from 25% to 75%, thus producing constraint trees *TB*; hence, *T*_25_ contains the 25% longest branches and *T*_75_ contains the 75% longest branches (and is therefore more resolved than *T*_25_).

For the biological dataset we used a constraint tree containing the “Magnificent Seven” (clades that were considered highly likely to be true, as these were the clades defining the strict consensus of the 2015 Prum et al. [12] tree and the 2014 Jarvis et al. [11] tree. On this dataset we also computed a tree using default ASTRAL (version 5.7.7).

Our evaluation criteria for the simulated datasets is the RF error rate for the estimated species tree. For the avian biological dataset since the true tree is not known, we evaluated results using MQSST scores and branch support (as calculated using ASTRAL’s local posterior probability values [29]). We also report wall clock running time for all analyses, noting that analyses of the simulated datasets were limited to four hours on the campus cluster.

Analyses of the simulated datasets were performed on the Illinois campus cluster, a heterogeneous computing cluster with a four-hour time limit. The analyses of the Jarvis *et al*. dataset with 48 species and 14,446 genes, required a more substantial computational environment; these were was run on the Tallis cluster on the Illinois campus cluster, which allows four weeks of analysis. Each task had at least 64GB of memory available. We run all methods single-threaded and use the recommended default settings when possible. For full details (locations of datasets, version numbers and commands, etc., see the supplementary materials at http://tandy.cs.illinois.edu/baqiao-suppl.pdf.

## 6 Results

### Results on simulated datasets

We show tree error rates across all model conditions in Figure 2. Error rates are lower when the constraint tree is better resolved (e.g., lower for *T*_75_ than for *T*_25_) and also lower when the number of genes increases; both these trends are expected. Error rates are also highest under the highest ILS condition (MC1, where AD=69%). For the avian simulated datasets, the error rates are higher with shorter species tree branches (i.e., avian 0.5X) than for the longer species tree branches (i.e., avian 2X), which is also the same as saying the error rates are higher under the higher ILS conditions. This trend is also expected.

**Figure 2:**
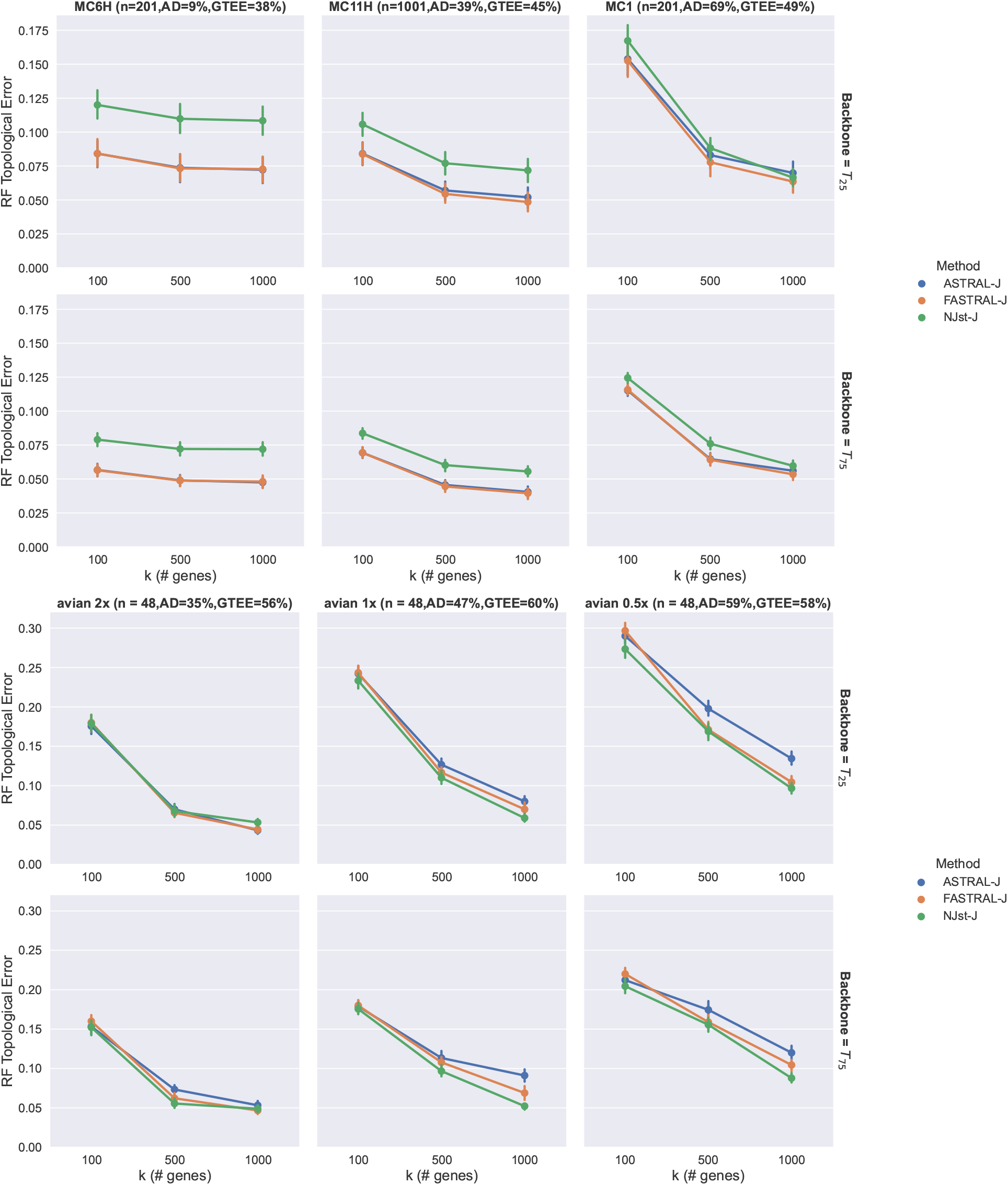
Tree error rates under the different model conditions with different constraint trees. Top two rows are for MC6H, MC11H, and MC1 model conditions, bottom two rows are for the avian simulated datasets with different branch lengths (0.5X, 1X, and 2X). For each model condition we explore two different constraint trees: *T*_25_ and *T*_75_, where *T*_*z*_ denotes the true species tree contracting all branches except the longest *z*% branches. The *x*-axis varies in *k* (number of genes). The RF topological error shown is the mean across 50 replicates under MC6H and MC1. Under MC11H, when backbone is *T*_25_, ASTRAL-J timed out on 16 replicates, and thus for MC11H the average on the completed 34 replicates is shown for all the methods. The error bars indicate standard error.

Other trends, and in particular the relative accuracy of the different methods, are less predictable. For example, the gap between methods in some cases decreases as the number of genes increase, but not in all cases. Interestingly, NJst-J is best under the avian simulated datasets but least accurate on the other model conditions. The relative performance between ASTRAL-J and FASTRAL-J is also somewhat variable. There is a large differences in accuracy between the two methods on the avian simulated datasets that favors FASTRAL-J, but little difference on the other model conditions.

The methods are, however, clearly differentiated in terms of running time (Figure 3). Across all conditions, NJst-J is the fastest of the three methods, and its running time is not impacted by the number of genes; this is a trend easily explained by the two-step process it uses where it first computes the internode distance matrix and then computes a tree from that distance matrix (only the first step is impacted by the number of genes, but it is a very fast step). The running time advantage of NJst-J over the other two methods depends on the model condition (amount of ILS, GTEE, and numbr of genes) as well as on the resolution of the constraint tree, so that the gap can be small given a highly resolved constraint tree (i.e., *T*_75_) and otherwise can be large. The largest gap on these simulated data was for the model condition MC11H, with 1001 species and 1000 genes; given the constraint tree *T*_25_ those datasets, ASTRAL-J used 9440 seconds (i.e., more than 2.6 hours), FASTRAL-J used about 1110 seconds (i.e., approximately 18.5 minutes), and NJst-J used about 139 seconds (i.e., just over 2 minutes).

**Figure 3:**
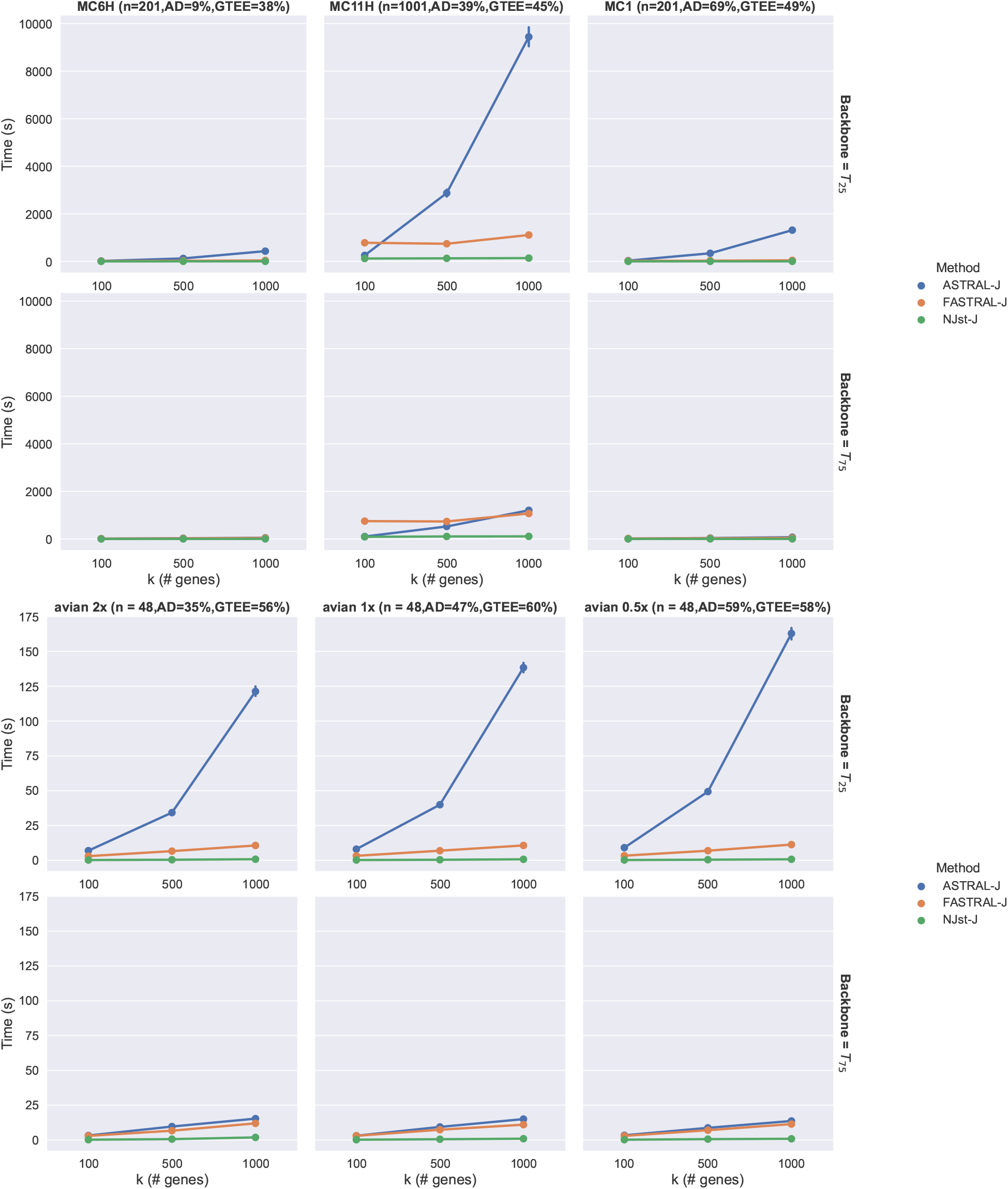
Wall-clock running times: top two rows are MC6H, MC11H, and MC1 datasets, and bottom two rows are avian simulated datasets. The running time shown is the mean across 50 replicates under MC6H and MC1. Under MC11H, when backbone is *T*_25_, ASTRAL-J timed out on 16 replicates, and thus for MC11H the average on the completed 34 replicates is shown for all the methods. The error bars indicate standard error.

**Figure 4:**
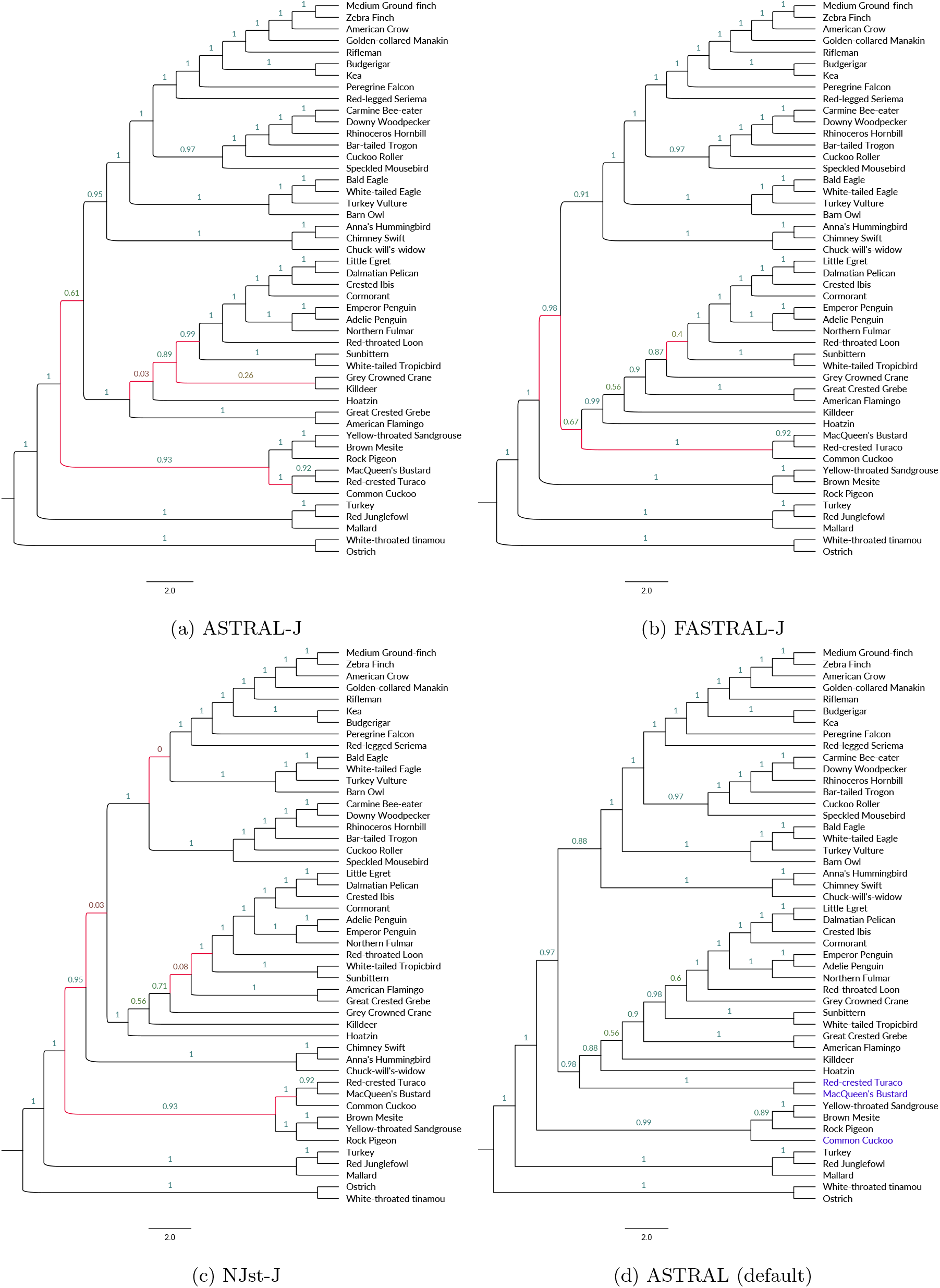
Results on the Jarvis et al. avian phylogenomic dataset [11] with 48 species and 14,446 genes estimated using ASTRAL-J, FASTRAL-J, NJst-J, and default ASTRAL (i.e., without a constraint tree). The constraint tree contains the “Magnificent Seven” clades suggested by [13]. Numbers denote localPP branch support. Red edges indicate differences to the default ASTRAL tree. Default ASTRAL violates one of the seven clades (clade IV, Ortidimorphae) by separating the three taxa (shown in blue).

The comparison for running time between FASTRAL-J and ASTRAL-J shows that in some cases they have very close running times, especially when the constraint tree is highly resolved. However, when there was a noteworthy difference between FASTRAL-J and ASTRAL-J, FASTRAL-J was faster. Furthermore, there are some cases where ASTRAL-J failed to complete analyses within the allowed four hour time limit on the Campus Cluster, but no other method failed in that time period. Thus, ASTRAL-J has a more substantial computational profile compared to the other methods.

### Results on the Avian Phylogenomic dataset of Jarvis *et al*

**[11]** In 2014, Jarvis et al. [11] published an avian phylogeny using 14,446 genes on 48 taxa. Their published tree, called the TENT (total evidence nucleotide tree) tree, was based on a RAxML concatenation analysis; among its major conclusions is that Neoaves splits into two clades, Columbea and Passerea. Several findings of the study, including the split of Neoaves into these two clades, are contested in a 2015 study by Prum et al. [12]. Reddy et al. [13] established a set of seven clades they considered highly reliable, because they were found in both the Prum and Jarvis trees, and referred to these as the “Magnificent Seven”. Subsequently, Braun and Kimball [31] reconsidered the Magnificent Seven and found that one of these clades (Clade IV, Otidimorphae) was not quite as clearly reliable as the others.

We used the Magnificent Seven to define a constraint tree (with only these seven clades). We then analyzed the 14,446 gene trees from [11] using ASTRAL-J, FASTRAL-J, and NJst-J with this constraint tree; we also computed the ASTRAL tree (run in default mode) on this dataset. Since the true tree is not known (beyond what is in the constraint tree), we evaluate the results using MQSST scores and branch support values (using local posterior probability); further, we report the running time of these methods on this dataset (Table 2).

**Table 2:**
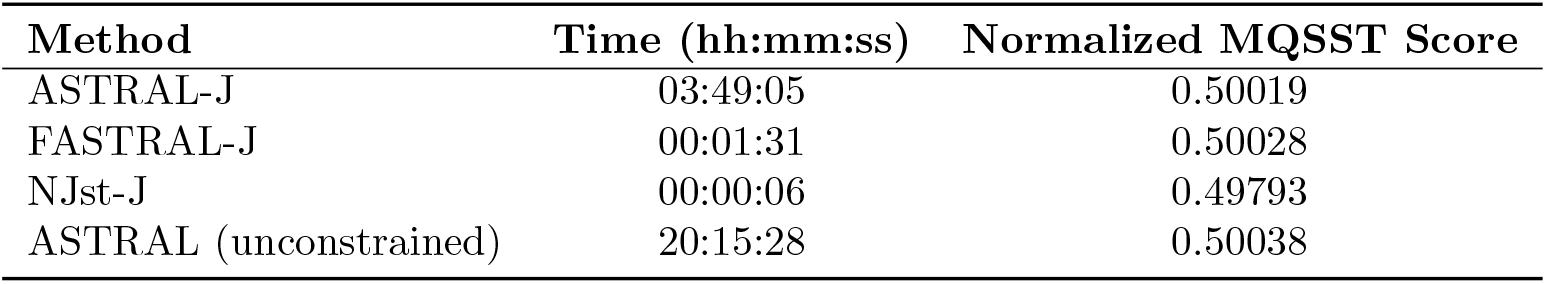
Running time (hours, minutes, and seconds) and normalized MQSST score of ASTRAL-J, FASTRAL-J, NJst-J, and defau;t ASTRAL on the avian biological dataset of [11] with 14,446 genes and 48 species. The constraint tree contains the “Magnificent Seven” clades [13].

| Method | Time (hh:mm:ss) | Normalized MQSST Score |
| --- | --- | --- |
| ASTRAL-J | 03:49:05 | 0.50019 |
| FASTRAL-J | 00:01:31 | 0.50028 |
| NJst-J | 00:00:06 | 0.49793 |
| ASTRAL (unconstrained) | 20:15:28 | 0.50038 |

ASTRAL run without any constraint tree satisfies six of the seven clades in the constraint tree, but violates Clade IV (Otidimorphae); interestingly, this is the only clade from the Magnificent Seven that was identified recently in [31] as not being sufficiently reliable to be considered a valid constraint. This analysis thus could be considered potential confirmation of the reliability of the remaining six clades, which were returned by ASTRAL when run without any constraints. This ASTRAL tree also has high MQSST score (0.50038) and good branch support: most branches are above 0.97, the lowest branch support in the tree is 0.56, and there are only two branches with support less than 0.75.

A comparison between the three tree-constrained analyses is also interesting. Their overall MQSST scores range from 0.49793 (for NJst-J) to 0.50028 (for FASTRAL-J), with ASTRAL-J in the middle at 0.50019. For the most part, all these trees are in topological agreement with each other (i.e., never differing in more than 7 out of 45 internal edges). However, although most branches are highly supported in all three trees, the trees do differ in terms of branch support. In particular, the NJst-J tree has three branches with support less than 70% (one with 56% support, one with 8%, and one with 3%). The ASTRAL-J tree has three branches with support less than 70%: one with 61%, one with 26%, and one with 3%, making it somewhat better supported than the NJst-J tree. The FASTRAL-J tree is somewhat better supported than the ASTRAL-J tree, with only three branches with support less than 70% (one with 67%, one with 56%, and one with 40%). Thus, of these three trees, FASTRAL-J has the highest branch support and the highest MQSST scores, ASTRAL-J is in second place, and NJst-J is in third place.

An interesting result of this study is how none of the trees contain the Columbea or Passerea clades; the monophyly of these two clades are two of the key differences between the Jarvis *et al*. tree (which found these to be monophyletic) and the Prum *et al*. tree (which did not find them to be monophyletic).

## 7 Discussion

We have presented FASTRAL-J and NJst-J, two species tree methods that allow users to provide unrooted non-binary trees as constraints on the final tree. Previous to this study, ASTRAL-J was the only summary method that allowed such constraints to be provided. Our simulation study shows that NJst-J is very fast, much faster than the other methods we explored, and that FASTRAL-J is also faster than ASTRAL-J except when the number of genes is very small. We also saw that FASTRAL-J either matched or improved on the accuracy of ASTRAL-J, and that NJst-J varied in its relative accuracy compared to ASTRAL-J (sometimes more accurate, sometimes less accurate). Overall, FASTRAL-J is generally the most accurate or close to the most accurate and is much faster than ASTRAL-J, while NJst-J (although the fastest) is not as reliable. Results on the avian phylogenomics dataset [11] are consistent with these trends: NJst-J is clearly the fastest (6 seconds), followed by FASTRAL-J (1.5 minutes), and then by ASTRAL-J (3 hours and 49 minutes). In addition, FASTRAL-J improved on both ASTRAL-J and NJst-J with respect to MQSST scores and branch support, showing its value on this biological dataset.

A major value in these tree-constrained methods is their greatly reduced speed compared to default ASTRAL. For example, default ASTRAL is dramatically slower on the avian phylogenomic dataset than all the other methods: it uses more than 20 hours and the slowest of the alternative methods uses less than 4 hours. Overall, while FASTRAL-J is particularly useful, all three tree-constrained species tree estimation methods provide benefits to biological systematics. In addition to speeding up species tree estimation, they enable biologists to explore the impact of assumptions about reliable clades; thus, multiple different hypotheses could be explored without the need for extensive computational resources.

